# MraZ is a transcriptional inhibitor of cell division in *Bacillus subtilis*

**DOI:** 10.1101/2022.02.09.479790

**Authors:** Maria L. White, Abigail Hough-Neidig, Sebastian J. Khan, Prahathees J. Eswara

## Abstract

The bacterial division and cell wall (*dcw*) cluster is a highly conserved region of the genome which encodes several essential cell division factors including the central divisome protein FtsZ. Understanding the regulation of this region is key to our overall understanding of the division process. *mraZ* is found at the 5’ end of the *dcw* cluster and previous studies have described MraZ as a sequence-specific DNA binding protein. In this article, we investigate MraZ to elucidate its role in *Bacillus subtilis*. Through our investigation, we demonstrate that increased levels of MraZ result in lethal filamentation due to repression of its own operon (*mraZ*-*mraW*-*ftsL*-*pbpB*). We observe rescue of filamentation upon decoupling *ftsL* expression, but not other genes in the operon, from MraZ control. Furthermore, through timelapse microscopy we were able to identify that overexpression of *mraZ*, results in de-condensation of the FtsZ ring (Z-ring). This is likely due to depletion of FtsL, and thus, we believe the precise role of FtsL is likely in Z-ring maturation and promotion of subsequent treadmilling. Our data suggests that regulation of the *mra* operon may be an alternative way for cells to quickly arrest cytokinesis potentially during entry into stationary phase and in the event of DNA replication arrest.

## Introduction

Bacterial cell division is a highly orchestrated process that typically results in the creation of two identical daughter cells through binary fission (1-3). Many species of bacteria encode a conserved gene neighbourhood known as the division cell wall (*dcw*) cluster (4, 5). In general, the peptidoglycan biosynthesis genes are found towards the 5’ end and genes encoding important cell division factors, including *ftsZ*, are found at the 3’ end (5-7). Gram-positive bacteria have an additional conserved region – the *ylm* operon, downstream of *ftsZ* which includes genes for additional cell division factors such as *sepF* and *divIVA* (8, 9).

At the very 5’ end of the *dcw* cluster is a gene encoding for a DNA binding protein MraZ (previously known as *yllB*), which is conserved in diverse lineages. In a range of species, including *Escherichia coli* and many Firmicutes *mraZ* is found within a short operon consisting of itself, *mraW* (*rsmH*; *yllC*), *ftsL* (*yllD*) and *pbpB (pbp2B)* (10). In genome-reduced Mycoplasma species, the *dcw* cluster consists of *mraZ and mraW* followed by *ftsA and ftsZ* alone (11).

Previous work in *E. coli* and Mycoplasma showed that overexpression of *mraZ* results in a change in transcriptional regulation of its own operon causing lethal filamentation in *E. coli* and cell enlargement in Mycoplasma (11-13). In *E. coli*, this phenotype could be resolved by co-expression of the gene immediately downstream of *mraZ, mraW*, a putative 16S rRNA (and possibly DNA) methyltransferase (12, 14). Work in *Burkholderia cenocepacia* has shown that *P*_*mra*_ is the sole transcription start site of the *dcw* cluster and MraZ can bind to the promoter sequence and presumably act as a transcriptional regulator (15). Recent investigation of Neisseriaceae family organisms revealed that deletion of *mraZ* among others factors may have allowed for the evolution of alternate growth modes (16). A recent report in *Staphylococcus aureus* proposes a role for MraZ in virulence regulation (17).

In this report, we show that overexpression of *mraZ* is toxic in *B. subtilis* due to cell division inhibition similar to what has been reported in other organisms thus far. Through fluorescence microscopy, we show that MraZ is a DNA-associated protein. Using transcriptional reporter assays and RNA-seq analysis, we elucidate that MraZ functions as a transcriptional repressor of the *mra* operon. Additionally, we demonstrate that MraZ-mediated lethal cell division inhibition is driven primarily by the reduction in the levels of FtsL, a critical divisome component that is turned over rapidly (18). Finally, we provide evidence that coalescence during maturation of FtsZ ring assembly is impaired upon MraZ overproduction presumably due to insufficient FtsL. Thus, our results together with studies conducted in other bacteria, favour the notion that MraZ is a transcriptional inhibitor of cell division.

## Results

### Overproduction of MraZ is lethal to *B. subtilis* and is dependent on DNA binding

To investigate the role of *mraZ* in *B. subtilis*, we constructed an IPTG-inducible copy of *mraZ* at an ectopic locus. We grew cultures of wildtype *B. subtilis* (WT) and cells containing inducible *mraZ* and plated serial dilutions on LB (lysogeny broth) agar with and without 1 mM IPTG. We found that when grown on IPTG, cells containing inducible *mraZ* (*mraZ*^*+*^) were unable to grow at any dilution in contrast to the WT control (**Fig. 1A**). This is similar to what has previously been shown in *E. coli* (12). Additionally, we investigated whether overexpression of *mraZ* resulted in a growth defect in liquid medium. After 3 h of 1 mM IPTG addition, cells overproducing MraZ display a drop in cell density at OD_600_ indicating cell lysis (**Fig. 1B**). To study the cause of lethality, we observed *mraZ*^*+*^ cells under the microscope. When grown in the presence of 1 mM IPTG for 2 h, *mraZ*^*+*^ cells are extremely filamentous (14.6 μm ± 6.9 μm) in contrast to the WT control (3.4 μm ± 1 μm) (**Figs. 1C and 1D**). This phenotype is indicative of cell division arrest in rod-shaped organisms. Eventually, the *mraZ* overexpressing cells go on to lyse explaining the lethal phenotype observed on solid and liquid media.

**Figure 1.**
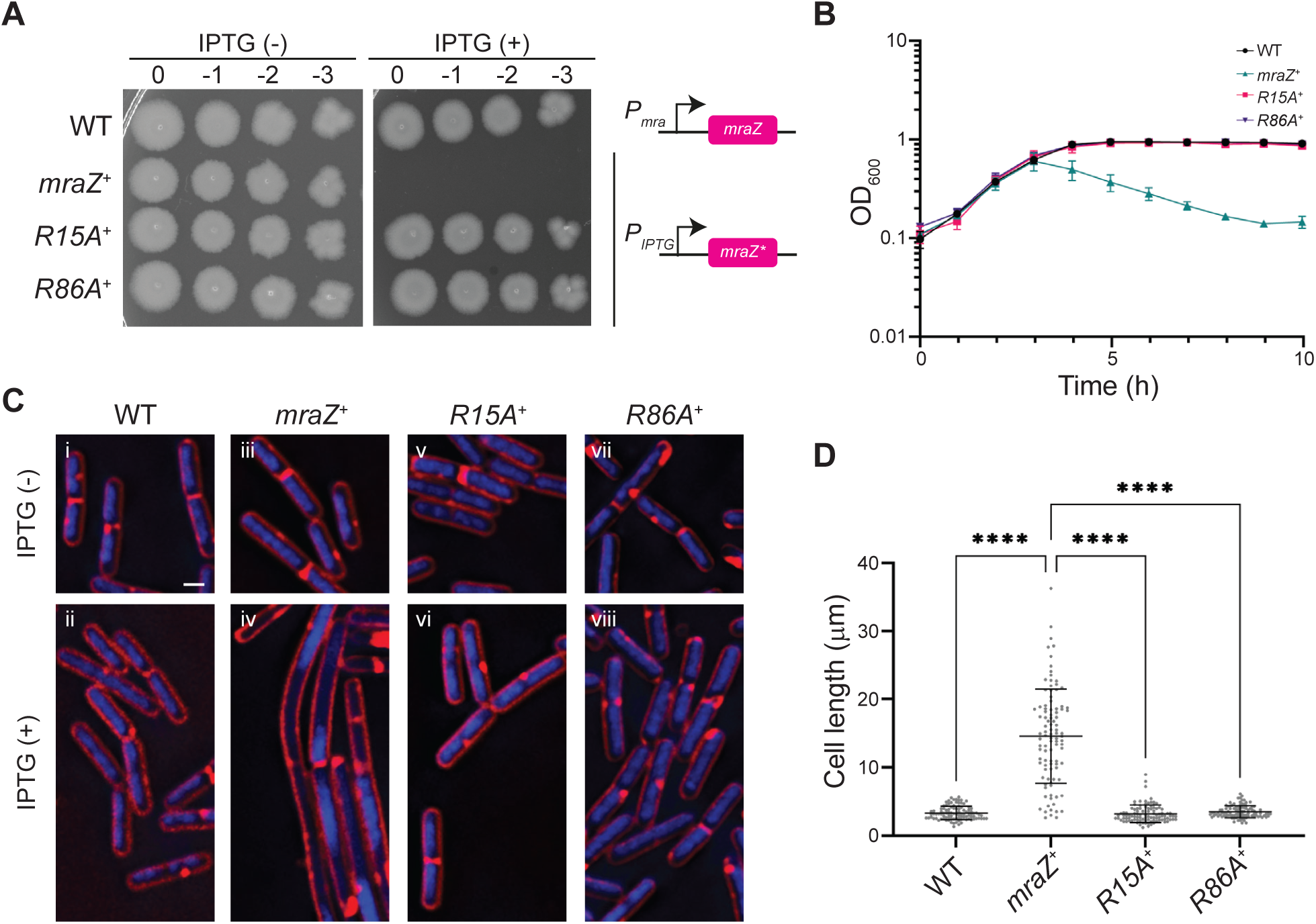
Overproduction of MraZ is lethal to *B. subtilis* and is dependent on DNA binding. (**A**) Spot assay of WT (PY79) *B. subtilis* and cultures containing inducible *mraZ*^*+*^ (MW189), *mraZ*^*R15A*^ (MW256) or *mraZ*^*R86A*^ (MW350), serially diluted and spotted onto agar in the absence and presence of 1 mM IPTG. Cartoon on the right depict the control of *mraZ*/mutants (*mraZ*\*) under native promoter or IPTG-inducible promoter. (**B**) Growth curve of WT *mraZ*^*+*^, *mraZ*^*R15A*^ and *mraZ*^*R86A*^ in the presence of 1 mM IPTG. Readings taken every hour at OD_600nm_. (**C**) Fluorescence micrographs of WT *B. subtilis* (i-ii), *mraZ*^*+*^ (iii-iv), *mraZ*^*R15A*^ (v-vi) and *mraZ*^*R86A*^ (vii-viii) in the presence and absence of 1 mM IPTG. Cell membrane stained with Synapto Red (FM4-64) and DNA stained with DAPI. Scale bar is 1 μm. (**D**) Quantification of cell length from microscopy in panel C. n = 100 cells; **** P < 0.0001.

We also noticed that MraZ overproducing cells exhibited severe impairment in the number of copies of chromosomes per cell suggesting a possible arrest in DNA replication. This is not due to filamentation by itself, as other conditions that elicit filamentation in *B. subtilis* such as the overexpression of *S. aureus gpsB* leads to no such DNA phenotype (19). This extreme nucleoid phenotype only occurs when MraZ is present in excess in the cell. At a lower IPTG concentration range, overproduction of MraZ slows the growth of the cells (**Fig. S1C**), produces filamentation, and some condensation of the nucleoid is observed but phenotypically the DNA is much more WT-like than at higher concentrations of IPTG (**Fig. S1B**).

MraZ belongs to the AbrB and SpoVT family of transcription factors and contains two highly conserved DXXXR motifs (**Fig. S1A**) (20-22). Previous work by Eraso *et al*. (12) had shown that a single point mutation of the first motif from arginine to alanine (R15A) was sufficient to prevent lethality in *E. coli*. We sought to identify whether this was the case for *B. subtilis* MraZ and generated point mutations in both DXXXR motifs – R15A and R86A. As described earlier, we serially diluted and plated both the R15A and R86A variants on LB agar with and without 1 mM IPTG. When grown in the presence of IPTG both mutants showed no lethality and grew similarly to the WT *B. subtilis* control (**Fig. 1A**). This was also seen in liquid culture (**Fig. 1B**). When imaged under the microscope both the R15A and R86A mutants were observed to be phenotypically WT-like in length (3.3 μm ± 1.28 μm and 3.5 μm ± 0.9 μm, respectively) in contrast to the filamentation observed in the unmutated *mraZ* overexpression strain (**Figs. 1C and 1D)**. Furthermore, there was no nucleoid phenotype when either of the two DXXXR mutants were overproduced. Thus, the ability of MraZ to cause filamentation, and thereby induce lethality relies on the presence of both of these motifs.

### MraZ associates with the chromosome through DXXXR motifs

To monitor whether MraZ localises to the chromosome we constructed a C-terminal fusion of MraZ to the green florescent protein (GFP). When overexpressed on solid media *mraZ-gfp* results in the formation of translucent colonies (**Fig. 2A**), however there is no significant growth defect in liquid media (**Fig. 2C**). This phenotype is in contrast to the overproduction of untagged MraZ which is toxic on both solid and liquid media. Overproduction of MraZ-GFP results in cells that are slightly shorter (10.4 μm ± 6.7 μm) than overproduction of untagged MraZ (15.7 μm ± 6.9 μm) but are nonetheless filamentous in comparison to the WT control (3.7 μm ± 1.1 μm), (**Figs. 2B and 2D**). However, in these cells the nucleoid appears WT-like in contrast to overproduction of untagged MraZ at 1 mM IPTG (**Fig. 1C**). GFP signal can be seen exclusively at the chromosome, similar to DNA-specific DAPI stain, indicating that MraZ associates with the nucleoid (**Fig. 2B**) (as noted previously (23)). Thus, it appears that MraZ has two functions, one that is responsible for filamentation and another for DNA replication inhibition, and addition of GFP tag to the C-terminus separates the two functions. The coating of entire nucleoid suggests that MraZ may play a larger role in nucleoid organization and/or as a transcriptional factor with control of many genes spread throughout the genome.

**Figure 2.**
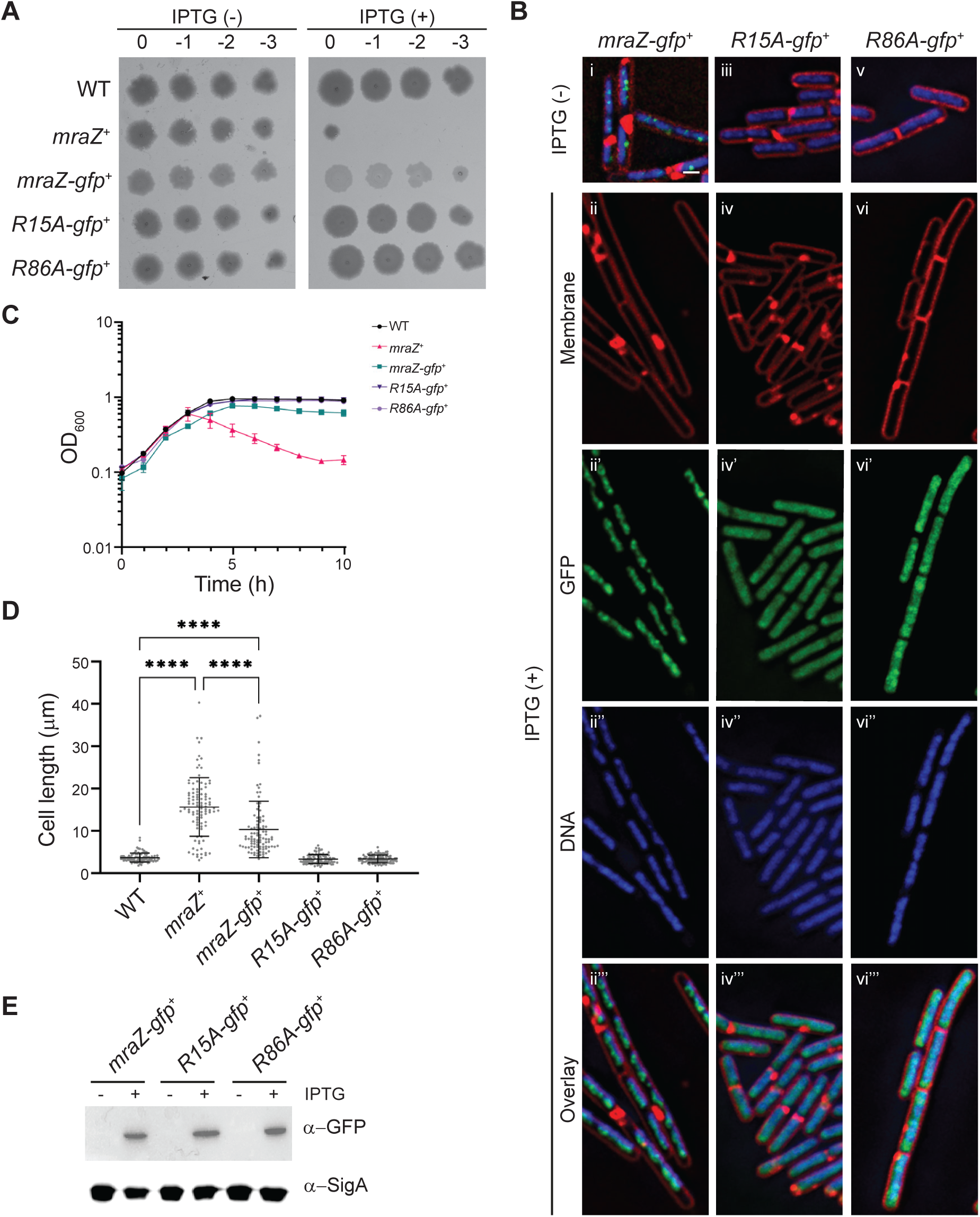
MraZ associates with the chromosome through DXXXR motifs. (**A**) Spot assay of cultures containing WT (PY79) *B. subtilis, mraZ*^*+*^ (MW189). *mraZ-gfp* (MW295), *mraZ*^*R15A*^*-gfp* (MW296) and *mraZ*^*R86A*^*-gfp* (MW351) serially diluted and spotted onto plates with and without 1 mM IPTG. (**B**) Micrographs of cells containing *mraZ-gfp* (i-ii), *mraZ*^*R15A*^*-gfp (*iii-iv) and *mraZ*^*R86A*^*-gfp* (v-vi) in the presence and absence of IPTG. Membrane is visualised with Synapto Red and DNA is stained with DAPI. Scale bar is 1μm. (**C**) Growth curve of WT, *mraZ*^*+*^, *mraZ-gfp, mraZ*^*R15A*^*-gfp* and *mraZ*^*R86A*^*-gfp* cultures following the addition of 1 mM IPTG, readings taken every hour at OD_600nm_. (**D**) Quantification of cell length in panel B. n = 100 cells, **** P < 0.0001. (**E**) Stability of MraZ-GFP and its mutants were confirmed through immunoblotting. Cell lysates from induced and uninduced cultures were probed with anti-GFP and anti-Sigma A (loading control) antibodies.

We tagged *mraZ*^*R15A*^ and *mraZ*^*R86A*^ to *gfp* to generate C-terminal GFP fusions to identify whether mutation of the DXXXR binding motif prevents MraZ from co-localising with the DNA, and in both cases GFP signal can be seen diffused in the cytoplasm, the cells are WT-like in length (R15A: 3.4 μm ± 1.1 μm, R86A: 3.4 μm ± 0.9 μm) and the nucleoids appear normal (**Figs. 2B and 2D**). We confirmed the stable production of tagged proteins via western blot (**Fig. 2E**). Indicating that diffuse signal is a result of MraZ-GFP mislocalisation and MraZ binding to the chromosome is dependent on the presence of both DXXXR DNA binding motifs.

### MraZ represses expression of the *mra* operon through the MraZ Binding Repeats (MBRs)

The promoter of the *mra* operon contains multiple MraZ binding repeats (MBRs) in a diverse range of species, including Mycoplasma, *E. coli* and *B. subtilis* (11). In *B. subtilis*, this repeat consists of three GTGG[A/T]G motifs separated by a 4-nucleotide spacer ((11); **Fig. 3A**). We were able to observe similar patterns in 7 other species that belong to the Firmicutes phylum and generated a sequence logo of the consensus sequence showing conservation of the GTGG repeat in the upstream region of the *mra* operon (**Fig. S2A**). Following a BLAST search of the PY79 genome, only one double or triple repeat was found within the promoter region of *mraZ*. In addition, we conducted a search of the PY79 genome through Pattern Locator (24) and were only able to identify a triple repeat in the upstream region of the *mra* operon confirming what was found through BLAST. Probing for only double repeats in Pattern Locator resulted in 13 hits (**Table S3**). Besides MBRs upstream of *mraZ*, only two hits landed in an intergenic region - one partially overlapping with the open reading frames of genes *mraY* and *murD* (part of the *dcw* cluster) and the other overlapping *yaaL* and the intergenic region upstream of *bofA*. A search using GTGGAG or GTGGTG single repeat resulted in 464 and 332 hits respectively. However, whether MraZ binds at these sites remains to be investigated.

**Figure 3.**
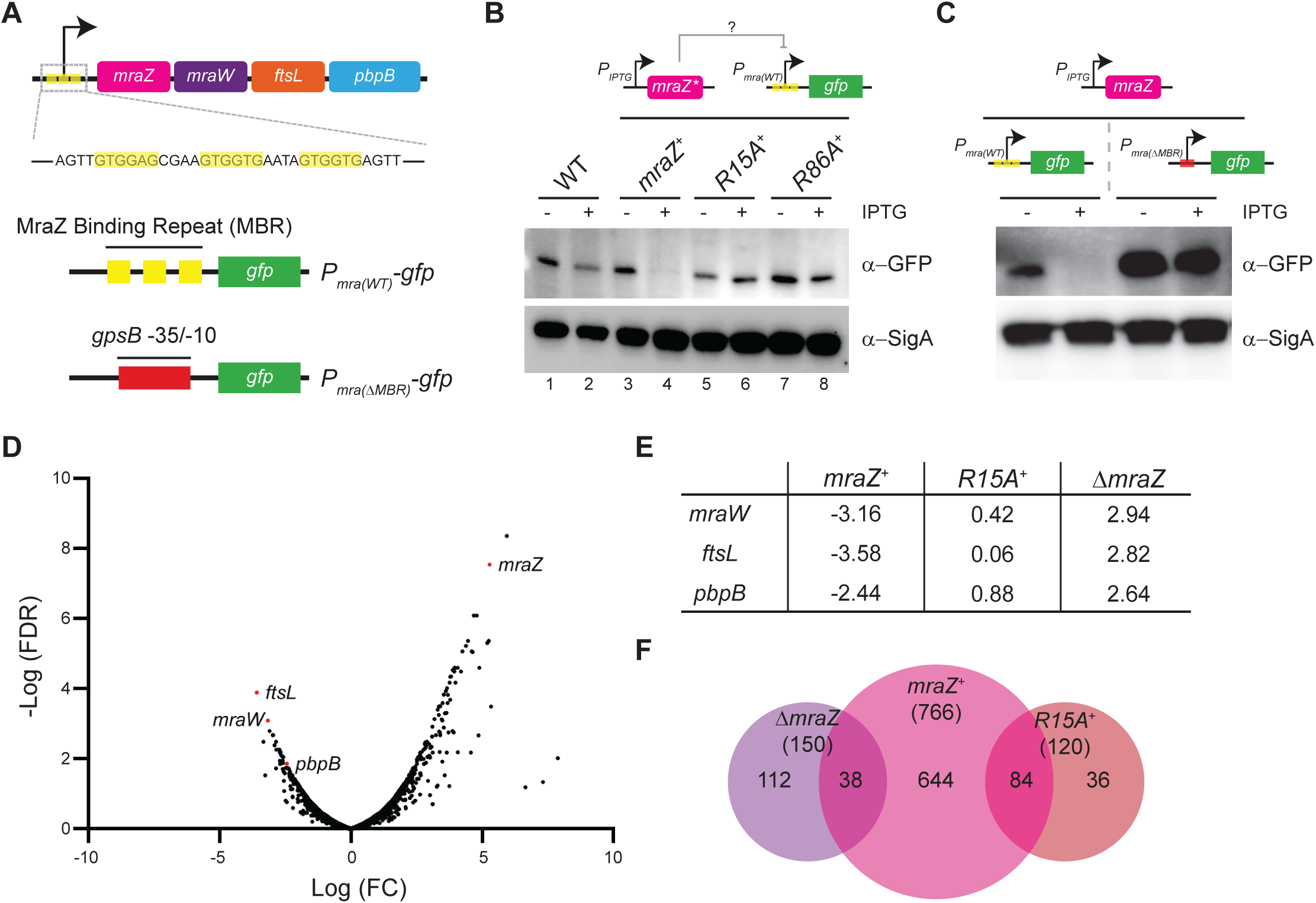
MraZ represses expression of the *mra* operon through the MraZ Binding Repeats (MBRs). (**A**) Cartoon representation of the *mra* promoter region and operon showing the binding repeats in yellow and the nucleotide sequence of the region below. Graphical representation of the transcriptional fusion of *P*_*mra*_ to *gfp* (MW368) and of the mutant *mra* promoter which has the MBRs removed and switched with the -35/-10 region of *gpsB* (MW478). (**B**) Immunoblot of cultures containing a transcriptional fusion of *P*_*mra*_ to *gfp* (MW368 – Lanes 1-2) with either *mraZ*^*+*^ (MW385 – Lanes 3-4), *mraZ*^*R15A*^ (MW389 – Lanes 5-6) or *mraZ*^*R86A*^ (MW429 – Lanes 7-8) in the presence and absence of 1 mM IPTG. Whole cell lysates were probed with anti-GFP and anti-Sigma A (loading control) antibodies. (**C**) Immunoblot of cultures containing either the wildtype *mra* promoter (MW385 – Lanes 1-2) or a mutant promoter lacking the MBRs (MW478 – Lanes 3-4) fused to GFP, with and without *mraZ* overproduction. Whole cell lysates were probed with anti-GFP and anti-Sigma A antibodies. (**D**) Volcano plot of RNA-Seq analysis from overexpression of *mraZ* (MW189). Points in red are genes within the *mra* operon. (**E**) Table showing the log fold change of RNA levels for the mra operon genes in *mraZ*^+^ (MW189), *mraZ*^*R15A*+^ (MW256) and Δ*mraZ* (MW192) compared to WT (PY79) control. (**F**) Venn diagram showing genes differentially regulated as identified through RNA-Seq when *mraZ* (MW189), *mraZ*^*R15A*^ (MW256) were overexpressed or *mraZ* was deleted (MW192) relative to WT (PY79). Total number of differentially regulated genes is indicated within the parenthesis.

To investigate the regulation of *mra* operon in *B. subtilis* we constructed a GFP-based transcriptional reporter of the *mraZ* promoter that includes all three MBR repeats (*P*_*mra*_*-gfp*) (**Fig. 3A**). This construct was introduced in the WT and *mraZ*^*+*^ backgrounds and the cell lysates were western blotted and probed with anti-GFP and anti-SigA antibodies (**Fig. 3B**). We observed a single band corresponding to the size of GFP from cultures containing *P*_*mra*_*-gfp* indicating the native MraZ control of the transcriptional reporter (**Fig. 3B**; lanes 1 and 2). In *mraZ*^*+*^ background we could detect GFP in the minus inducer control, however in the plus inducer condition GFP was below the detectable range (**Fig. 3B**; lanes 3 and 4). This indicates that overproduction of MraZ leads to strong repression of the *mra* promoter. We next tested whether the overproduction of either of the DXXXR mutants (R15A or R86A) disrupted the ability to repress the promoter of *mraZ*. Our results indicate that neither R15A nor R86A is able to repress *P*_*mra*_*-gfp* (**Fig. 3B**; lanes 5-8). Thus, MraZ-mediated repression of the *mra* promoter is dependent on its ability to bind DNA through both DXXXR motifs.

We were interested in determining whether the repression of MraZ on the *mra* promoter was dependent or independent of the MBRs. To this end we constructed a variation of the fusion in which the -35/-10 region of *P*_*mra*_ was switched with that of *gpsB* – a constitutively expressed cell division gene that is not part of the *dcw* cluster (**Fig. 3A**). We introduced each of these fusions into WT or *mraZ*^*+*^ strain backgrounds. In these backgrounds we then overexpressed *mraZ* and western blotted the resulting cultures. We found that overexpression of *mraZ* did not result in repression of *P*_*mra*_ in the absence of MBRs (**Fig. 3C**). Therefore, these results confirm that the MBRs are a requirement for MraZ mediated repression of the *mra* operon.

While the triple repeat is only found within *P*_*mra*_, GTGG[A/T]G is a common sequence within the genome and we sought to identify whether MraZ transcriptionally regulates any other genes in *B. subtilis* through RNA-Seq analysis. We were particularly interested as MraZ-GFP appears to bind promiscuously throughout the chromosome (**Fig. 2B**). RNA was extracted from *B. subtilis* cells overproducing MraZ, MraZ^R15A^ and additionally from cells lacking *mraZ*. We identified 766 differentially regulated genes when *mraZ* was overexpressed (**Fig. 3F**), through comparison of the dataset obtained when the inactive *mraZ*^*R15A*^ was overproduced we were able to eliminate some of these non-specific changes in gene expression (**Fig. 3F**). We then compared the remaining 682 differentially regulated genes against those that were up or down regulated in a Δ*mraZ* background. This resulted in a set of 38 genes, however on further analysis 34 of those 38 genes are upregulated when *mraZ* is overexpressed as well as when *mraZ* is deleted, making it unlikely that these genes are part of the MraZ regulon (**Table S1**). Three of the remaining 4 differentially regulated genes were part of the *mra* operon (*mraW, ftsL*, and *pbpB*) and are downregulated when *mraZ* is overexpressed and upregulated when *mraZ* is deleted (**Figs. 3D and 3E**). Additionally, we observed that *mraZ* overexpression results in a decrease in the mRNA level of *pksJ*, which encodes for a polyketide synthetase, and in Δ*mraZ* strain level of *pksJ* mRNA is increased (**Supplemental File 1**). However, whether MraZ is directly regulating this gene and the physiological reasons for the change in expression under these conditions is yet to be elucidated. Overall, based on our analysis it appears that MraZ primarily acts as an autoregulatory transcriptional repressor of the four-gene operon (*mraZ*-*mraW*-*ftsL*-*pbpB*). It is possible that association at other sites may facilitate nucleoid organization.

### Repression of *ftsL* by MraZ drives cell division arrest

Two of the 4 genes in the *mra* operon (*ftsL* and *pbpB*) are known to be essential in *B. subtilis* (10, 25). As the operon includes two essential cell division genes, we sought to identify which genes were responsible for the filamentation we observed when MraZ was overproduced. To that end, we uncoupled the expression of *mraW, ftsL*, and *pbpB* from MraZ control by placing each of these genes individually under an inducible promoter. We then expressed *mraZ* with each one of the other genes in the operon to determine if there was any resolution to the filamentation we previously observed. We first confirmed that *mra* repression is occurring at all IPTG concentrations tested to identify the lowest concentration that resulted in filamentation (**Figs. S1B-S1E**). Based on our results, we chose 25 μM IPTG for this set of experiments.

When *mraW* is uncoupled from MraZ repression, cells are still filamentous (11.6 μm ± 5.8μm) and a similar length to overexpression of *mraZ* alone (10.8 μm ± 6.0μm) (**Figs. 4A and 4B**). This result is in contrast to what was previously observed in *E. coli* where co-expression of *mraZ* and *mraW* results in a rescue of the lethal phenotype (12). When *pbpB* is overexpressed with *mraZ* the cells are again filamentous and similar in length to *mraZ* overexpression (12.0 μm ± 6.0 μm). Thus, suggesting that although *pbpB* is an essential gene, immediate cell division arrest may not be due to repression of its expression, as PBP2B appears to be stable (26). In contrast, when we uncoupled expression of *ftsL* from MraZ control, we observed cells that were much shorter (5.7 μm ± 2.1 μm) and closer to wildtype in length (3.9 μm ± 1.0 μm). Thus, it appears that the repression of *ftsL* is the likely cause of immediate cell division arrest in MraZ overproducing cells, as FtsL is rapidly turned over (18, 26-29).

**Figure 4.**
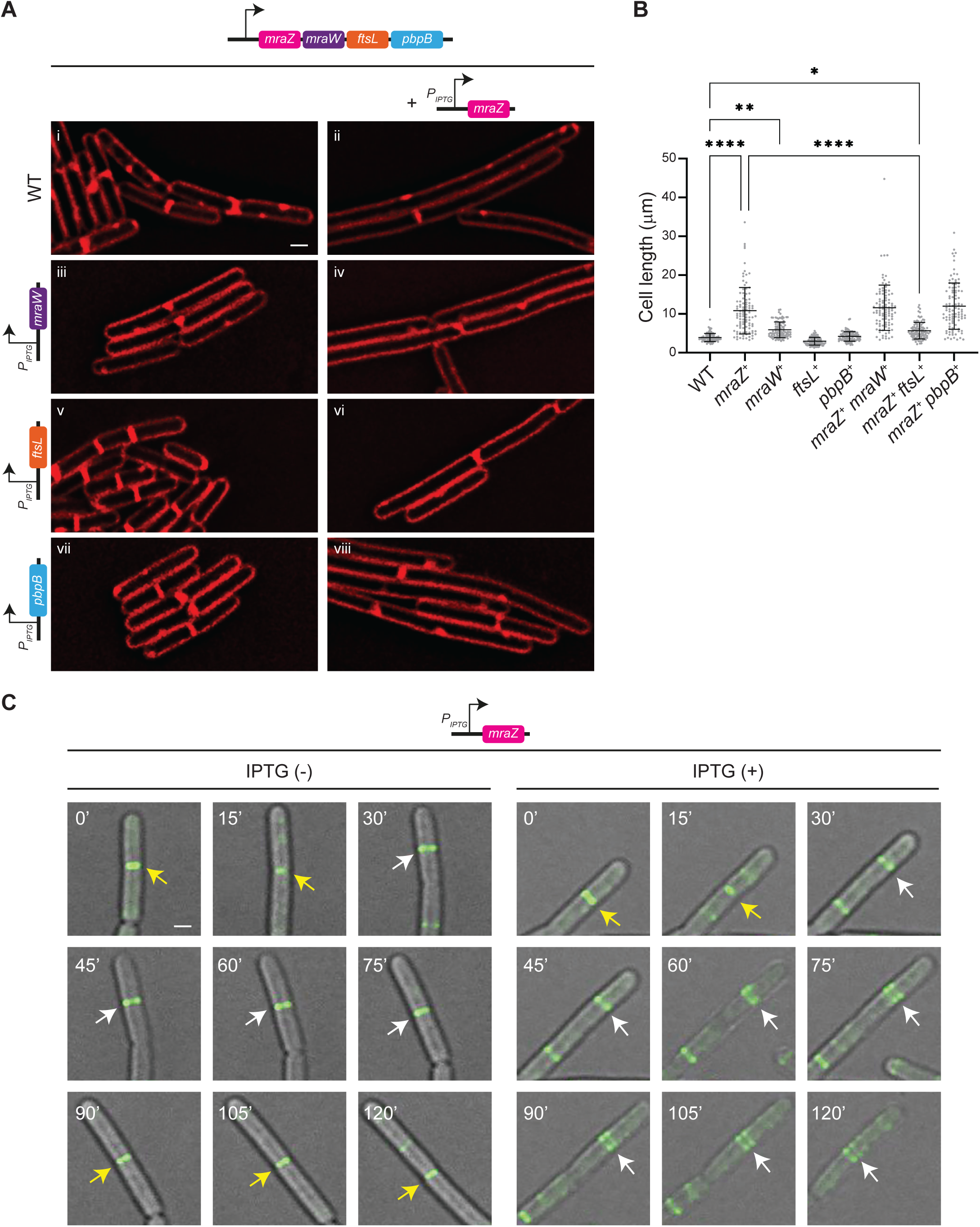
Repression of *ftsL* by MraZ drives cell division arrest. (**A**) Fluorescence micrographs of WT (PY79) *B. subtilis* (i) and cells expressing *mraZ*^*+*^ (MW289) (ii), *mraW*^*+*^ (MW379) (iii), *ftsL*^*+*^ (MW330) (v), or *pbpB*^*+*^ (BAH1) (vii); and of cells co-expressing *mraZ*^*+*^*/mraW*^*+*^ (MW387) (iv), *mraZ*^*+*^*/ftsL*^*+*^ (MW333) (vi) and *mraZ*^*+*^*/ pbpB*^*+*^ (BAH4) (viii) in the presence of 25 μM IPTG. Membrane is visualised with Synapto Red, scale bar is 1 μm. (**B**) Quantification of cell length from microscopy in panel B. n=100; **** P < 0.0001, ** P = 0.06, * P = 0.0264 (**C**) Timelapse microscopy of cultures containing *mraZ*^*+*^ harboring *ftsZ-gfp* (MW205) in the absence and presence of 1 mM IPTG. Images were taken every 15 minutes for 2 hours. Yellow arrows indicate constriction-efficient FtsZ rings and white arrows follow newly assembled FtsZ rings. Scale bar is 1 μm.

### MraZ-mediated repression of *ftsL* results in de-condensation of the FtsZ ring

Knowing that decoupling of *ftsL* expression relieves filamentation when MraZ is overproduced, we wanted to study the immediate effects of FtsL depletion on divisome assembly. To investigate this, we utilised a strain containing *ftsZ-gfp* (30), to track FtsZ dynamics when MraZ is in excess. Using timelapse microscopy we monitored the localisation pattern of FtsZ-GFP at 15-min intervals in the *mraZ*^+^ strain in the absence and presence of IPTG. In the absence of IPTG, cells progress through the division cycle normally (**Fig. 4C**; **Movie 1**). However, after the addition of IPTG, cells can be observed dividing regularly with the Z-ring forming and constricting initially. After 60 mins, Z-rings that have formed but have not constricted begin to de-condense. By 75 mins after induction, multiple Z-rings can be seen immediately adjacent to one another (**Fig. 4C**; **Movie 2)**, suggesting a likely impairment of the formation of functional Z-rings that are capable of undergoing constriction. Given that decoupled expression of *ftsL* from MraZ control resolves filamentation (**Figs. 4A and 4B**), we speculate that the main role of FtsL is in the formation of constriction-capable Z-ring assembly.

Additionally, we investigated the effect of deleting *mraZ* on cell division, as in this strain background expression of *ftsL* (and *pbpB*) is unregulated. Upon imaging cells lacking *mraZ*, we observed that on average these cells are smaller than WT and appear very similar to cells overproducing FtsL (**Figs. S2B and S2C**), indicating hyperactivation of cytokinesis. We did not observe similar short-cell phenotype in *pbpB*^+^ strain (**Fig. 4AB**). These results further provide evidence that the principal role of FtsL is in Z-ring maturation and constriction. This phenotype has previously been observed following deletion of the RasP protease which is known to facilitate FtsL turnover (28).

## Discussion

Cell division is a highly orchestrated and complex process, the timing of which must be tightly regulated. Numerous signals feed into the decision to divide including, population density, nutrient availability and the status of the chromosome. Previous reports have shown that MraZ is an important transcriptional regulator, indeed in some species such as *E. coli*, MraZ regulates at least the first 9 genes in the *dcw* cluster and may be responsible for controlling transcription as far downstream as FtsZ (5, 31-33). Among the Gram-positive Actinobacteria phylum, work in *Corynebacterium glutamicum* has shown that MraZ mRNA is degraded by RNase III and that loss of *rnc* leads to accumulation of *mraZ* mRNA, and therefore increased cellular levels of MraZ. This results in cell elongation through MraZ repression of *ftsEX*, however whether MraZ in *C. glutamicum* is a transcriptional repressor of the *mra* operon is yet to be elucidated (34). Upon search of *B. subtilis* MraZ binding motifs in other organisms, we noted the high level of conservation in species within and outside of the Firmicute phylum (**Fig. S2A and Table S5**).

Whilst the structure and function of MraZ and its multimeric crystal structure are fairly well characterised at least in certain species (12, 35, 36), the role of its syntenous partner, MraW, remains unclear. MraW is predicted to be a 16S rRNA methyltransferase (N^4^ cytosine C1402). It has been reported that chloroplast MraW and 16S rRNA methylation may play a role in ribosome levels (37). Work in *E. coli* suggests that MraW may be able to regulate codon utilization (38), and possibly act as a transcriptional regulator through methylation of DNA (14). Additionally results in *S. aureus* suggests that MraW may play a role in virulence (39). However, whilst *E. coli* MraZ and MraW appear to be regulated by each other we were unable to identify a similar relationship in *B. subtilis* (12). Curiously, through our RNA-Seq analysis we were able to identify an upregulation of genes encoding ribosomal proteins when *mraZ* is deleted - conditions under which levels of MraW are likely increased (**Table S4**). However, the biological relevance of this change remains to be elucidated.

In this report, we show that similar to previous studies conducted in other organisms, MraZ is a transcriptional regulator in *B. subtilis* (11-13, 16). Specifically, it is important in repressing the expression of two essential cell division genes *pbpB* and *ftsL*, in addition to the non-essential *mraW* gene described above, which does not appear to have a direct role in cell division (10) (**Fig. 5A**). Although the essential nature of FtsL has been well characterized, the precise role of FtsL in the divisome complex remains unclear in *B. subtilis*. Studies in *E. coli* have elucidated that FtsL is involved in initiating membrane invagination and septal peptidoglycan synthesis that accompanies Z-ring treadmilling under the direction of FtsN (40-43). However, FtsN is absent in *B. subtilis* (44). As shown in **Fig. 5B**, in *B. subtilis*, FtsL forms a complex with PBP2B (class B PBP; *E. coli* PBP3/FtsI homolog) and DivIC (FtsL paralog; *E. coli* FtsB homolog), and it is known that PBP2B also interacts with DivIB (*E. coli* FtsQ homolog) and FtsW (shape, elongation, division and sporulation (SEDS) family glycosyltransferase) (45-51). It is noteworthy that although PBP2B is essential, its catalytic function is not (52). Thus, the essentiality of PBP2B comes from its scaffolding role. Based on our studies, we reveal that the FtsL level is integral for constriction-efficient FtsZ ring assembly (**Fig. 5C**). Upon MraZ overproduction (in the absence of enough FtsL), we show that the Z-rings that have coalesced are unable to retain the structure and disassemble into multiple Z-rings. This is reminiscent of what was noted when FtsZ associated proteins (ZAPs) were absent (53). Similarly, defective Z-ring assembly has been noted when FtsW/PBP1 (homolog of *B. subtilis* PBP2B) levels are synthetically lowered in *S. aureus* (54). Perhaps the role of FtsL is to stabilize the FtsW/PBP2B complex and is conserved across multiple species. Absence of EzrA (one of the FtsZ anchors) in a *B. subtilis* strain producing less FtsL is lethal, and overexpression of *ftsL* restores delayed FtsZ ring constriction in cells lacking *ezrA* (55). Thus, FtsL complex may communicate with ZAPs and transduce that signal to kickstart the constriction/FtsZ treadmilling process, similar to what has been reported in *E. coli* (40-42, 56). However, how the activation of FtsZ treadmilling is triggered by optimal level of FtsL in the absence of FtsN-like proteins remains to be investigated.

**Figure 5.**
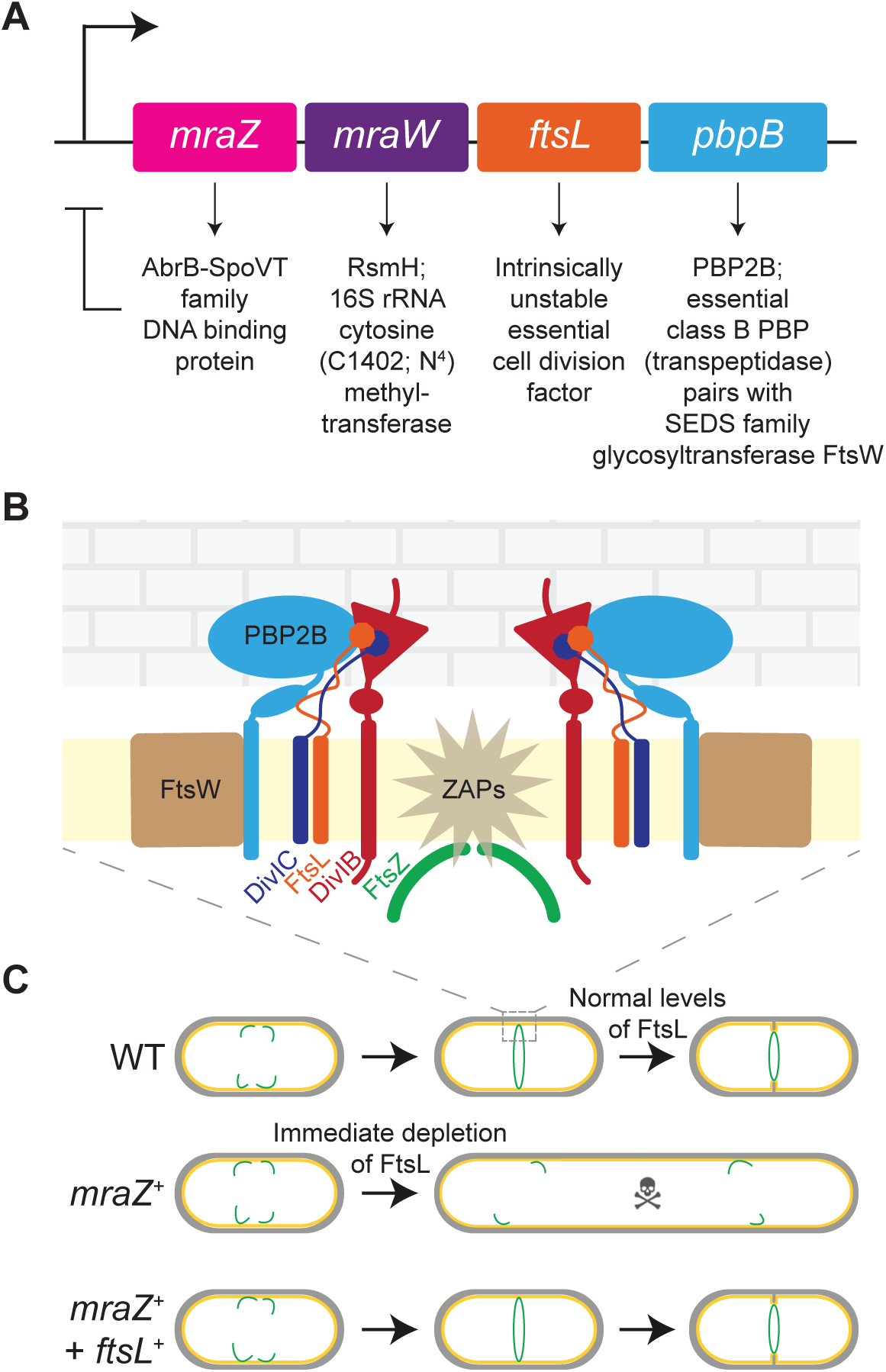
Cell division inhibition by MraZ is mediated primarily through FtsL. Genes of the *mra* operon and the respective functions of their protein products. FtsL is an intrinsically unstable protein that serves as a linchpin for the constriction-efficient divisome assembly. To achieve this FtsL directly interacts with PBP2B (of the FtsW/PBP2B - SEDS/class B PBP complex), DivIC, DivIB, and perhaps also some FtsZ-associated proteins (ZAPs) either directly or indirectly. **(C)** Lethal phenotype of *mraZ* overexpression stems (at least initially) from the depletion of FtsL, which results in the lack of mature constriction-capable FtsZ (shown in green) ring formation. Additional supply of *ftsL* results in restoration of cell length that is similar to that of WT.

Among the divisome components, FtsL has emerged as a key factor to regulate cell division in a rapid manner (27, 28). FtsL is a transmembrane protein that is an essential part of the divisome, loss of which causes extreme filamentation in both *E. coli* and *B. subtilis* (29, 57). Previous studies have shown that FtsL is intrinsically unstable (58) and is rapidly turned over by the membrane protease RasP (18, 27, 28, 59). At elevated growth temperature, DivIB has been shown to protect FtsL from rapid degradation (18); and a strong complex between DivIC and FtsL has also been alluded to aid in the stabilization of FtsL (27, 45, 46). It was previously suggested that accumulation of FtsL at the divisome may be a rate determining step (28). The results we report here, further validates the previous findings that the level of FtsL is critical for cell division. This unstable nature of FtsL, combined with its essentiality makes it an attractive control point to arrest cell division. In fact, during DNA-damage response the known cell division inhibitor, YneA, appears to halt division by interacting with late arriving divisomal proteins including FtsL and Pbp2B in *B. subtilis* (60). Interestingly, YneA mediated cell division inhibition could be rescued by *ftsL* overexpression (55). Similarly, another study discovered that during phage SP01 infection, a phage protein gp56 inhibits *B. subtilis* cell division by possibly directly interacting with FtsL (61). In this report, we show how in addition to post-translational regulation of FtsL to regulate cell division, transcriptional regulation of *mra* operon by MraZ could also be an efficient way to halt cell division due to the intrinsic instability of FtsL. Indeed, it has been proposed that the *mra* operon may be directly regulated by DNA replication initiation protein, DnaA, upon inhibition of DNA replication (62). In further support of this notion of transcriptional inhibition of cell division, it appears that during entry into stationary phase the transcriptional activity of *mraZ* significantly increases and that of the genes in rest of the operon *mraW, ftsL*, and *pbpB* drops, presumably to eventually halt division (10) (**Fig. S2D**).

## Materials and methods

### Strain Construction and General Methods

All *B. subtilis* strains utilised in this study are derivatives of PY79 (63) and were constructed via double recombination of circular plasmids into the chromosome. Information regarding the strains used can be found in **Table S1**. All plasmids were constructed using standard cloning protocols (PCR amplification, restriction digest-ligation). Plasmids were transformed into the parental strain directly and integration confirmed via screening of the *amyE* locus for pDR111 (D. Rudner) based plasmids or the *thrC* locus for pDG1664 based plasmids. For generation of strains that require two separate genes of interest to be integrated into the *amyE* locus, a background containing a second synthetic *amyE* locus was used (AHB286: *bkdB::TB917::amyE::catR;* A. Camp).

#### mraZ^+^

*mraZ* was amplified from the PY79 chromosome using primer pair oMW114/oMW115 (See **Table S2** for oligonucleotide details), the resulting fragment was then digested with HindIII and NheI and ligated into pDR111; an IPTG inducible vector for integration at the *amyE* locus in *B. subtilis*. The resulting plasmid pMW35 was then transformed into *B. subtilis* producing MW189.

#### mraZ^R15A^/mraZ^R86A^

*mraZ* was amplified with oMW114/oMW115 from a geneblock (Integrated DNA Technologies) that contained either the R15A (gMW4) or R86A (gMW5) mutation, before being digested and ligated into pDR111 as for MW189. See **Table S2** for the geneblock sequence. The plasmids pMW49 (R15A) and pMW65 (R86A) were then transformed into PY79 *B. subtilis* producing strains MW256 and MW350 respectively.

#### mraZ-gfp

*mraZ* was amplified with oMW114/oMW172, the resulting product was then digested with HindIII and BamHI, additionally *linker-gfp* was amplified from geneblock gP1 with primers oMW173/oMW121, the fragment was then digested with BamHI and NheI before being ligated into pDR111. This generated plasmid pMW59 which was transformed into *B. subtilis* generating strain MW295. Tagged point mutants (*mraZ*^*R15A*^*-gfp* and *mraZ*^*R86A*^*-gfp)* were constructed as above with *mraZ* amplified from gMW4 or gMW5. The resulting plasmids pMW58 (R15A) and pMW66 (R86A) were then used to transform *B. subtilis* generating strains MW296 and MW351 respectively.

#### P_mraZ_-gfp

Promoter fusions of *mraZ* were constructed in the *thrC* integration plasmid pDG1664 (64). The promoter region of *mraZ* (500 bp upstream of the *mraZ* start codon) was amplified with oMW177/oMW178 from chromosomal DNA and *gfp* with oMW184/oMW185. The resulting fragments were digested with BamHI/XhoI and XhoI/EcoR1 respectively and then ligated into pDG1664, producing plasmid pMW68. Another fusion that lacked the MraZ binding repeats was made by amplifying the region containing the sequence upstream of the MBRs with oMW177/oMW252, this was then digested with BamHI/XhoI and ligated with *gfp* that was amplified with two rounds of PCR with oMW253 (oMW254-R2)/oMW185 to add on the -35/-10 region from *gpsB* and digested with XhoI/EcoRI, which was then ligated into pDG1664 producing plasmid pMW81. Both plasmids were transformed into PY79 generating MW368 and MW452 respectively, pMW35 was then used to transform these strains generating MW385 and MW478. Additionally, pMW49 and pMW65 were used to transform MW368 resulting in MW389 and MW429 respectively.

#### ftsL^+^/mraW^+^/pbpB^+^

Inducible *ftsL, mraW and pbpB* were constructed by amplifying the respective genes from PY79 chromosomal DNA with primer pairs, oMW127/oMW128, oMW122/oMW123 and oMW275/oMW276, the resulting fragments were digested with SalI and NheI before being ligated into pDR111. This created plasmids pMW39 (*ftsL*^*+*^), pMW38 (*mraW*^*+*^) and pAH1(*pbpB*^*+*^), which were then used to transform AHB286, which contains a second synthetic *amyE* locus, producing strains MW330, MW379 and BAH1 respectively. To construct strains that additionally have inducible *mraZ*, MW189 was transformed with pDAG32 (65) to switch the resistance cassette to *catR* generating strain MW289. The chromosomal DNA from this strain was then used to transform MW330, MW379 and BAH1, resulting in strains MW387, MW333 and BAH4 respectively. pMW39 was additionally used to transform PY79 to produce MW207.

#### Δ*mraZ*

Chromosomal DNA from the Bacillus Genetic Stock centre (BGSC) strain BKK15130 (66) was used to transform PY79 to generate MW192 (*mraZ::kanR*).

#### mraZ^+^ ftsZ-gfp

AD3007 (30), a strain which has an additional copy of *ftsZ-gfp* at the native locus was transformed with pMW35 resulting in strain MW205.

### Media and Culture Conditions

Unless otherwise stated, overnight cultures of *B. subtilis* strains were grown at 22 °C in lysogeny broth (LB) and diluted 1:10 in fresh LB before being grown at 37 °C to mid-log phase (OD_600nm_= 0.5-0.8) cultures were then standardised to OD_600nm_= 0.1 in fresh LB. Where induction of genes under IPTG required, unless otherwise stated, a final concentration of 1 mM IPTG was added when cultures were standardised, and the cultures were then grown for 2 h at 37 °C.

### Spot Assays

All spot assays were carried out similarly to previously described (67). Briefly strains were grown in liquid culture at 37 °C whilst shaking until mid-log phase (OD_600nm_ = 0.5-0.8) before being standardised to OD_600nm_=0.1. Standardised cultures were then serially diluted before 1 μl of each serial dilution was spotted onto either lysogeny agar (LA) or LA supplemented with 1 mM IPTG when required to induce expression of genes under IPTG control. Plates were incubated for approximately 14 h at 37 °C before being observed for any growth defects.

### Growth Curves

Strains to be analysed by growth curve were grown to mid-log phase (OD_600nm_ = 0.5-0.8) in liquid culture at 37 °C with agitation and were then standardised to OD_600nm_ = 0.1 in LB. IPTG was added to a final concentration of 1 mM where required. An aliquot of 250 μl of each standardised culture was added in triplicate to a 96 well plate, which was then incubated at 37 °C with agitation for 12 h in a Synergy H1 Plate Reader Gen. 5 (BioTek). OD_600nm_ readings were taken every hour and growth curves were plotted using GraphPad Prism 9.

### Microscopy

Microscopy was carried out as previously described (68). Briefly, 1 ml aliquots of *B. subtilis* cultures were pelleted and then washed with 1x phosphate buffer saline (PBS) via centrifugation. Pellets were then resuspended in 100 μl of PBS and were stained with Synapto Red to stain the membrane at a final concentration of 1 μg/ml and DAPI to stain the DNA – final concentration 1 μg/ml, 5 μl of the stained cells were spotted onto the base of a glass bottomed dish (MatTek) and 1% agarose pad made with PBS was placed gently on top. Imaging was carried out at room temperature using a DeltaVision Elite deconvolution fluorescence microscope with photos taken using a Photometrics CoolSnap HQ2 camera. All images were acquired by taking 17 z-stacks at 200-nm intervals. Images were deconvolved though the SoftWorx imaging software provided by the microscope manufacturer. Analysis of cell lengths was done through ImageJ with statistical analysis carried out using GraphPad Prism 9.

### Timelapse Microscopy

As described previously, strain MW205 was grown to mid-log phase (OD 0.5-0.8) before 5 μl was aliquoted onto a MatTek dish and covered with 1% agarose pad made with LB (68). Samples were allowed to adjust to the microscope chamber at 30 °C for a period of 30 min before 10 μl of 10 mM IPTG was added to the agarose pad for the plus inducer conditions. Timelapse imaging was immediately begun, and images were taken every 15 min for 2 h with 5 Z-stacks at 200 nm intervals. Image processing was carried out as described above.

### Immunoblots

Cultures were grown as previously described and 1 ml aliquots standardised to an OD_600nm_ of 1 were taken. Cells were pelleted and resuspended in 500 μl of protoplast buffer containing 0.5 M sucrose, 20 mM MgCl_2_, 10 mM KH_2_PO_4_ and 0.1 mg/ml lysozyme. Samples were then incubated for 30 min at 37 °C and prepared for SDS-PAGE. For analysis of *P*_*mra*_ activity across different growth phases, a 20 ml culture was standardised to OD_600nm_ of 0.1 and samples were taken every hour for 4 h, each sample was standardised to an OD_600nm_ of 1 in 1 ml and then prepared as previously described. Following electrophoresis samples were transferred to nitrocellulose membrane using the iBlot 2 transfer system (ThermoFisher) and probed with antibodies against GFP and *B. subtilis* SigA.

### Bioinformatics

Using the consensus sequence blastn was used to find MBRs in *B. subtilis* by utilizing the PY79 whole genome sequence (CP006881.1) as the query, and the search string: GTGGWGNNNNGTGGWGNNNNGTGGWG. Using algorithm parameters optimized for short nucleotide repeats with spaces (expect threshold 100000, word size 7, match/mismatch cost 1,-1, gap cost existence 0, extension 2, no filters or masks). To generate a sequence logo (https://weblogo.berkeley.edu/; (69)), sequence upstream of *mraZ* from the following species were used: *B. subtilis, Bacillus cereus, Staphylococcus aureus, Staphylococcus epidermidis, Streptococcus pneumoniae, Enterococcus faecium, Lactococcus lactis*, and *Listeria monocytogenes*. Visual examination of the sequences using the previously identified consensus sequence for *B. subtilis mraZ* binding site GTGG revealed the existence of ordered repeats containing the motif GTGGNNNNNAGTGGNGNNNNGTGG (11). The sequences can be found in **Table S5** and the multiple sequence alignment generated with the help of Clustal Omega (70) is shown in **Fig. S2A**. In addition, using the probe GTGGNNNNNNGTGG Pattern Locator (24) was used to search the PY79 genome for potential MraZ binding repeats (**Table S3**).

### RNA-Seq

Cultures were grown as previously described and treated with RNAprotect Bacteria (Qiagen) before RNA was extracted utilising the RNeasy Mini Kit (Qiagen). Samples were sent to Microbial Genome Sequence Centre (MiGS) for sequencing and analysis. GraphPad Prism 9 was used to generate a volcano plot.

## Acknowledgements

We thank our lab members for comments on the manuscript. This work was funded by the National Institutes of Health grant (R35GM133617) to PE.

## Author contributions

The conception and design of the study (MW, PE), data acquisition (MW, AH, SK, PE), analysis and/or interpretation of the data (MW, AH, SK, PE), and writing of the manuscript (MW, PE).

## Notes

### Competing Interest Statement

The authors have declared no competing interest.

