## Supplemental table and figures for "MraZ is a transcriptional inhibitor of cell division in *Bacillus subtilis*"

#### Supplemental figure, table, spreadsheet, and movie legends

**Figure S1**
(**A**) Multiple sequence alignment of the protein sequence of MraZ from *Bacillus subtilis*, *Staphylococcus aureus*, *Mycoplasma pneumoniae,* and *Escherichia coli.* Highlighted in green are the conserved DXXXR motifs, *, :, and ., indicate fully, strongly, or weakly conserved residues respectively. (**B**) Fluorescence micrographs of WT (PY79) *B. subtilis* and cells containing inducible *mraZ* at a range of IPTG concentrations. Membrane is visualised with Synapto Red and the DNA is stained with DAPI. Scale bar is 1 µm. (**C**) Growth curve of cells containing an IPTG inducible copy of *mraZ* (MW189) at 0 µM, 25 µM, 100 µM, 250 µM and 1 mM IPTG. $OD_{600nm}$ readings taken every hour. (**D**) Quantification of cell length from microscopy in panel B. n=100 **** P < 0.0001. (**E**) Immunoblot of cultures containing $P_{mra}$-*gfp* transcriptional reporter with *mraZ*$^+$ under control of an IPTG inducible promoter (MW385). Whole cell lysates were prepared from cultures grown without (0 µM) and with inducer (25 µM, 100 µM, 250 µM and 1000 µM IPTG) and were subsequently probed with anti-GFP and anti-Sigma A (loading control) antibodies.

**Figure S2**
(**A**) Sequence alignment and logo showing the conservation of the MraZ binding repeats (MBR) in different Firmicute species – *Bacillus subtilis*, *Bacillus cereus*, *Staphylococcus aureus*, *Staphylococcus epidermidis*, *Streptococcus pneumoniae*, *Enterococcus faecium*, *Lactococcus lactis*, and *Listeria monocytogenes*. * indicates a fully conserved nucleotide. (B) Fluorescence micrographs of WT (PY79) *B. subtilis*, Δ*mraZ* (MW192) and *ftsL*$^+$ (MW207) following the addition of 1 mM IPTG. Membrane is stained with Synapto Red. Scale bar is 1 µm. (C) Quantification of cell length from microscopy in panel A. n=200; **** P < 0.0001, ** P = 0.0022. (D) Immunoblot of cultures containing $P_{mra}$-*gfp* during different growth phases. Cultures were standardised to an $OD_{600nm}$, samples were taken every hour for 4 h and were subsequently probed with anti-GFP and anti-Sigma A (loading control) antibodies.

**Table S1** Strains used in this study.

**Table S2** Oligonucleotides and geneblocks.

**Table S3** GTGG double repeats in the PY79 genome.

**Table S4** Ribosome associated proteins are upregulated when mraZ is absent.

**Table S5** The MraZ Binding Repeats are conserved throughout the domains of life.

**Supplemental File 1** – Excel spreadsheet with the results of RNA-Seq analysis of *mraZ*$^{+}$ (MW189), *mraZ*$^{R15A}$ (MW256) or Δ*mraZ* (MW192). All values are relative to WT (PY79) *B. subtilis.* Results represent analysis of a pooled biological triplicate. Tabs indicate the genes that were identified as differentially expressed in each of the indicated backgrounds and additionally the expression levels of all identified genes in all three conditions. The 38 genes identified in **Fig. 3F** are listed.

**Movie 1 –** MW205 (*mraZ*$^{+}$ *ftsAZ:ftsZ-gfp*) grown in the absence of IPTG. Images taken every 15 min for 2 h. Scale bar is 2 μm.

**Movie 2 –** MW205 (*mraZ*$^{+}$ *ftsAZ:ftsZ-gfp*) grown in the presence of IPTG. Images taken every 15 min for 2 h following the addition of IPTG. Scale bar is 2 μm.

#### Table S1 - Strains used in this study

| Strain | Genotype | Reference |
|---|---|---|
| PY79 | Wildtype *B. subtilis* | (1) |
| AHB286 | bkdB::TB917::amyE::catR | Amy Camp |
| MW189 | amyE::$P_{hyperspank}$-mraZ$^{BS}$ specR | This study |
| MW256 | amyE::$P_{hyperspank}$-mraZ$^{R15A}$ specR | This study |
| MW350 | amyE::$P_{hyperspank}$-mraZ$^{R86A}$ specR | This study |
| MW295 | amyE::$P_{hyperspank}$-mraZ-linker-gfp specR | This study |
| MW296 | amyE::$P_{hyperspank}$-mraZ$^{R15A}$-linker-gfp specR | This study |
| MW351 | amyE::$P_{hyperspank}$-mraZ$^{R86A}$-linker-gfp specR | This study |
| MW368 | thrC::$P_{mraZ}$-gfp ermR | This study |
| MW385 | thrC::$P_{mraZ}$-gfp ermR amyE::$P_{hyperspank}$-mraZ$^{BS}$ specR | This study |
| MW389 | thrC::$P_{mraZ}$-gfp ermR amyE::$P_{hyperspank}$-mraZ$^{R15A}$ specR | This study |
| MW429 | thrC::$P_{mraZ}$-gfp ermR amyE::$P_{hyperspank}$-mraZ$^{R86A}$ specR | This study |
| MW478 | thrC::$P_{mraZ(-MBR)}$-gfp ermR amyE::$P_{hyperspank}$-mraZ$^{BS}$ specR | This study |
| MW379 | bkdB::TB917::amyE::$P_{hyperspank}$-mraW specR | This study |
| MW330 | bkdB::TB917::amyE::$P_{hyperspank}$-ftsL specR | This study |
| BAH1 | bkdB::TB917::amyE::$P_{hyperspank}$-pbpB specR | This study |
| MW289 | amyE::$P_{hyperspank}$-mraZ catR | This study |
| MW387 | bkdB::TB917::amyE::$P_{hyperspank}$-mraW specR amyE::$P_{hyperspank}$-mraZ catR | This study |
| MW333 | bkdB::TB917::amyE::$P_{hyperspank}$-ftsL specR amyE::$P_{hyperspank}$-mraZ catR | This study |
| BAH4 | bkdB::TB917::amyE::$P_{hyperspank}$-pbpB specR amyE::$P_{hyperspank}$-mraZ catR | This study |
| MW192 | mraZ::kanR | Derived from BGSC* BKK15130 (2) |
| MW207 | amyE::$P_{hyperspank}$-ftsL specR | This study |
| MW205 | amyE::Phyperspank-mraZ ftsAZ::ftsAZ-gfp Ω | This study; derived from (3) |

*BGSC: Bacillus Genetic Stock Center

#### Table S1 - References

1. Youngman P, Perkins JB, Losick R. 1984. Construction of a cloning site near one end of Tn917 into which foreign DNA may be inserted without affecting transposition in *Bacillus subtilis* or expression of the transposon-borne *erm* gene. Plasmid 12:1-9.
2. Koo BM, Kritikos G, Farelli JD, Todor H, Tong K, Kimsey H, Wapinski I, Galardini M, Cabal A, Peters JM, Hachmann AB, Rudner DZ, Allen KN, Typas A, Gross CA. 2017. Construction and Analysis of Two Genome-Scale Deletion Libraries for *Bacillus subtilis*. Cell Systems 4:291-305.e7.
3. Gregory JA, Becker EC, Pogliano K. 2008. *Bacillus subtilis* MinC destabilizes FtsZ-rings at new cell poles and contributes to the timing of cell division. Genes and Development 22:3475-3488.

### Table S2 - Oligonucleotides and geneblocks

Restriction sites are underlined and ribosome binding sites are in red for oligonucleotides. Mutated codons for gMW4 and gMW5 are in red, lower-case letters indicated mutated base pair. Linker sequence is indicated in red for gP1 and gP2.

| Primer | Sequence 5' to 3' |
|---|---|
| oMW114 | AATGA AAGCTT ACATAAGGAGGAACTACT ATGTTTATGGGTGAATATCAGCATACC |
| oMW115 | AATGA GCTAGC TTATATATCAAACCCAATCATGTTTTCAGCA |
| oMW172 | AATGA GGATCC TATATCAAACCCAATCATGTTTTCAGCAAT |
| oMW173 | AATGA GGATCC GGTTCCGCTGGCTCC |
| oMW121 | AATGA GCTAGC TTATTTGTATAGTTCATCCATGCCATGTG |
| oMW212 | AATGA AAGCTT ACATAAGGAGGAACTACT ATGAGTAAAGGAGAAGAACTTTTCACTG |
| oMW213 | AATGA GCTAGC TCCGGAACCTTCAAGGCCAGAAC |
| oMW214 | AATGA GCTAGC ATGTTTATGGGTGAATATCAGCATACC |
| oMW219 | AATGA GCATGC TTATATATCAAACCCAATCATGTTTTCAGCA |
| oMW122 | AATGA GTCGAC ACATAAGGAGGAACTACT ATGTTTCAACACAAGACAGTACTTCTT |
| oMW123 | AATGA GCTAGC TTATTTTCTTTTTTCAGCAATCCGAAGC |
| oMW127 | AATGA GTCGAC ACATAAGGAGGAACTACT ATGAGCAATTTAGCTTACCAACCAG |
| oMW128 | AATGA GCTAGC TCATTCCTGTATGTTTTTCACTTTTTTATCTTT |
| oMW275 | AATGA GTCGAC ACATAAGGAGGAACTACT ATGATTCAAATGCCAAAAAAGAATAAATT |
| oMW276 | AATGA GCTAGC TTAATCAGGATTTTTAAACTTAACCTTGA |
| oMW177 | AATGA GGATCC CGCCAGATTCCGGAA |
| oMW178 | AATGA CTCGAG TAACTCACCACTATTCACCACTTC |
| oMW184 | AATGA CTCGAG ACATAAGGAGGAACTACT ATGAGTAAAGGAGAA |
| oMW185 | AATGA GAATTC TTATTTGTATAGTTCATCCATGCCATGTGTAATC |
| oMW252 | AATGA CTCGAG AACTTCATTTTACACTGTTTACCCATAA |
| oMW253 | CAACCGTACAGATTTCTGAAAAAATATATGTGATGATTCACATAAGGAGGAACTA |
| oMW254 | AATGA CTCGAG GAGAGAGAAATTTGACAACCGTACAGA |
| oP4 | AAGTCTTATTTCCATAACTTTAGG |
| oP5 | GGCCAAAAAAACTGCTGCC |
| oP209 | TGGCAAGAACGTTGCTCGAGGGTAAATGTGAGCACTCAC |
| oP210 | CAATAATTGGTACGTACGATCTTTCAGCCGACTCAAACATC |

| Geneblock | Sequence 5' to 3' |
|---|---|
| gMW4 | ATGTTTATGGGTGAATATCAGCATACCATCGATGCGAAAGGCgcCATGATCGTACCCGCTAAATTCAGAGAAGGCCTA<br>GGTGAGCAATTTGTGCTGACTAGAGGACTTGACCAATGTCTCTTCGGCTACCCTATGCACGAATGGAAACAAATTGA<br>AGAAAAACTAAAAGCTCTTCCTCTCACAAAGAAAGATGCCCGCGCGTTTACCCGTTTCTTCTTTTCAGGGGCGACTGA<br>ATGCGAACTGGATAAGCAAGGCAGGGTAAATATCGCATCATCTCTATTGAATTACGCCAAACTGGAAAAAGAATGTGT<br>TGTTATCGGGGTTTCTAATCGAATTGAATTGTGGAGTAAAGTAATTTGGGAACAATACACAGAAGAGCAAGAAGATTC<br>ATTTGCTGAAATTGCTGAAAACATGATTGGGTTTGATATATAA |
| gMW5 | ATGTTTATGGGTGAATATCAGCATACCATCGATGCGAAAGGCCGCATGATCGTACCCGCTAAATTCAGAGAAGGCCT<br>AGGTGAGCAATTTGTGCTGACTAGAGGACTTGACCAATGTCTCTTCGGCTACCCTATGCACGAATGGAAACAAATTG<br>AAGAAAAACTAAAAGCTCTTCCTCTCACAAAGAAAGATGCCCGCGCGTTTACCCGTTTCTTCTTTTCAGGGGCGACTG<br>AATGCGAACTGGATAAGCAAGgcGCGGTAAATATCGCATCATCTCTATTGAATTACGCCAAACTGGAAAAAGAATGTG<br>TTGTTATCGGGGTTTCTAATCGAATTGAATTGTGGAGTAAAGTAATTTGGGAACAATACACAGAAGAGCAAGAAGATT<br>CATTTGCTGAAATTGCTGAAAACATGATTGGGTTTGATATATAA |
| gP1 | GGTTCCGCTGGCTCCGCTGCTGGTTCTGGCCTTGAAGGTTCCGGAATGAGTAAAGGAGAAGAACTTTTCACTGGAG<br>TTGTCCCAATTCTTGTTGAATTAGATGGTGATGTTAATGGGCACAAATTTTCTGTCAGTGGAGAGGGTGAAGGTGATG<br>CAACATACGGAAAACTTACCCTTAAATTTATTTGCACTACTGGAAAACTACCTGTTCCATGGCCAACACTTGTCACTAC<br>TTTCGCGTATGGTCTTCAATGCTTTGCGAGATACCCAGATCATATGAAACAGCATGACTTTTTCAAGAGTGCCATGCC<br>CGAAGGTTATGTACAGGAAAGAACTATATTTTTCAAAGATGACGGGAACTACAAGACACGTGCTGAAGTCAAGTTTGA<br>AGGTGATACCCTTGTTAATAGAATCGAGTTAAAAGGTATTGATTTTAAAGAAGATGGAAACATTCTTGGACACAAATTG<br>GAATACAACTATAACTCACACAATGTATACATCATGGCAGACAAACAAAAGAATGGAATCAAAGTTAACTTCAAAATTA<br>GACACAACATTGAAGATGGAAGCGTTCAACTAGCAGACCATTATCAACAAAATACTCCAATTGGCGATGGCCCTGTC<br>CTTTTACCAGACAACCATTACCTGTCCACACAATCTGCCCTTTCGAAAGATCCCAACGAAAAGAGAGACCACATGGTC<br>CTTCTTGAGTTTGTAACAGCTGCTGGGATTACACATGGCATGGATGAACTATACAAATAA |
| gP2 | ATGAGTAAAGGAGAAGAACTTTTCACTGGAGTTGTCCCAATTCTTGTTGAATTAGATGGTGATGTTAATGGGCACAAA<br>TTTTCTGTCAGTGGAGAGGGTGAAGGTGATGCAACATACGGAAAACTTACCCTTAAATTTATTTGCACTACTGGAAAA<br>CTACCTGTTCCATGGCCAACACTTGTCACTACTTTCGCGTATGGTCTTCAATGCTTTGCGAGATACCCAGATCATATG<br>AAACAGCATGACTTTTTCAAGAGTGCCATGCCCGAAGGTTATGTACAGGAAAGAACTATATTTTTCAAAGATGACGGG<br>AACTACAAGACACGTGCTGAAGTCAAGTTTGAAGGTGATACCCTTGTTAATAGAATCGAGTTAAAAGGTATTGATTTT<br>AAAGAAGATGGAAACATTCTTGGACACAAATTGGAATACAACTATAACTCACACAATGTATACATCATGGCAGACAAA<br>CAAAAGAATGGAATCAAAGTTAACTTCAAAATTAGACACAACATTGAAGATGGAAGCGTTCAACTAGCAGACCATTAT<br>CAACAAAATACTCCAATTGGCGATGGCCCTGTCCTTTTACCAGACAACCATTACCTGTCCACACAATCTGCCCTTTCG<br>AAAGATCCCAACGAAAAGAGAGACCACATGGTCCTTCTTGAGTTTGTAACAGCTGCTGGGATTACACATGGCATGGA<br>TGAACTATACAAAGGTTCCGCTGGCTCCGCTGCTGGTTCTGGCCTTGAAGGTTCCGGA |

#### Table S3 - GTGG double repeats in the PY79 genome

Using Patten Locator (https://www.cmbl.uga.edu/software/patloc.html) the sequence GTGGNNNNNNGTGG was probed for in the *B. subtilis* PY79 genome resulting in 13 hits, 9 of which were within the open reading frame (ORF) of the gene indicated (*). Two were part of the MraZ binding repeat upstream of *mraZ* (’). The remaining two were within the intergenic region of (or overlapping) the indicated genes (^). The pattern sequence and start/end position of the pattern in the genome is shown.

| # | Start | End | Pattern Sequence | Gene/proximal gene |
|---|---|---|---|---|
| *1^* | 29696 | 29709 | GTGGAAGTAAGTGG | *yaaL-bofA* |
| *2** | 517277 | 517290 | GTGGTTAATGGTGG | *yddR* |
| *3** | 711989 | 712002 | GTGGATGGGTGTGG | *yeeF* |
| *4** | 1206158 | 1206171 | GTGGAGGAATGTGG | *tenA* |
| *5** | 1257199 | 1257212 | GTGGCAATCGGTGG | *yjjA* |
| *6** | 1539312 | 1539325 | GTGGTTTCAGGTGG | *gerR* |
| *7’* | 1543103 | 1543116 | GTGGAGCGAAGTGG | *mraZ* |
| *8’* | 1543113 | 1543126 | GTGGTGAATAGTGG | *mraZ* |
| *9^* | 1551929 | 1551942 | GTGGTTATAAGTGG | *mraY-murD* |
| *10** | 2037173 | 2037186 | GTGGGACACAGTGG | *czrA* |
| *11** | 2057081 | 2057094 | GTGGAATGTCGTGG | *sodF* |
| *12** | 2499347 | 2499360 | GTGGAAGCCGGTGG | *yqbJ* |
| *13** | 3317493 | 3317506 | GTGGAATGCCGTGG | *rsbQ* |

#### Table S4 - Genes of ribosomal proteins are upregulated when *mraZ* is absent.

RNA-Seq analysis revealed that the genes encoding 9 ribosome proteins are upregulated when *mraZ* is deleted (MW192). Also indicated is the LogFC when *mraZ* is overexpressed (MW189) or when *mraZ*$^{R15A}$ (MW256) is overexpressed. All values are relative to WT (PY79) *B. subtilis.* Raw reads can be found in Supplemental File 1.

| | | Δ*mraZ* | | | *mraZ*$^{+}$ | | | *mraZ*$^{R15A}$ | | |
|---|---|---|---|---|---|---|---|---|---|---|
| Gene | Description | LogFC | P value | FDR | LogFC | P value | FDR | LogFC | P value | FDR |
| *rpsQ* | 30S ribosomal protein S17 | 1.69 | 0.01171 | 0.54186 | 0.28 | 0.46172 | 0.07414 | 0.46 | 0.48107 | 1 |
| *rpmC* | 50S ribosomal protein L29 | 1.67 | 0.01306 | 0.56501 | 0.35 | 0.52172 | 0.09865 | 0.52 | 0.42681 | 1 |
| *rplV* | 50S ribosomal protein L22 | 1.48 | 0.02621 | 0.93075 | 0.10 | 0.32618 | 0.02007 | 0.33 | 0.61821 | 1 |
| *rpsS* | 30S ribosomal protein S19 | 1.46 | 0.02825 | 0.96039 | -0.03 | 0.48504 | 0.00401 | 0.49 | 0.45921 | 1 |
| *rplW* | 50S ribosomal protein L23 | 1.45 | 0.02930 | 0.97950 | 0.11 | 0.71313 | 0.02084 | 0.71 | 0.27769 | 1 |
| *rplP* | 50S ribosomal protein L16 | 1.38 | 0.03830 | 1 | 0.73 | 0.14651 | 0.25170 | 0.15 | 0.82310 | 1 |
| *rplX* | 50S ribosomal protein L24 | 1.36 | 0.04022 | 1 | 0.15 | 0.23092 | 0.03271 | 0.23 | 0.72428 | 1 |
| *rplN* | 50S ribosomal protein L14 | 1.36 | 0.04087 | 1 | 0.65 | 0.15515 | 0.21999 | 0.16 | 0.81282 | 1 |
| *rpsC* | 30S ribosomal protein S3 | 1.32 | 0.04597 | 1 | 0.48 | 0.00331 | 0.14867 | 0.003 | 0.99613 | 1 |

#### Table S5 - The MraZ Binding Repeats are conserved throughout the domains of life.

Using BLAST search the sequence 100 bp upstream of the homologous mraZ coding site was identified in various species and their respective genome sources are indicated. Green indicates the conserved motif, yellow indicates the variable inter-motif sequence. Only members of the Firmicute phylum were used for WebLogo generation (**Fig. S2A**).

| Kingdom-Phylum | Species | Strain or Isolate | Accession | Location | Raw Sequence |
|---|---|---|---|---|---|
| Bacteria-Firmicutes | *Bacillus subtilis* | PY79 | CP006881.1 | 1543044-1543144 | TAAAGAACCCTGACTAGTTCAGGGTTTTTTTTTATGGGTAAACAGTGTAAAATGAAGTTGTGGAGCGAAGTGGTGAATAGTGGTGAGTTAAGGAGAGAAAG |
| Bacteria-Firmicutes | *Bacillus cereus* | DE0207 | NZ_VTQW01000024.1 | 51878-51978 | TGGACCGCCACCTCCCCTTTGTCGACCCTCGGGCGTCCCCATTCCCTACCTTGATGCCCCACGATACTCCACTTTGCTCCACCGTCAACGAAGTTTCGCCT |
| Bacteria-Firmicutes | *Listeria monocytogenes* | EGD-e | NC_003210.1 | 2125496-2125596 | TTAGACTTCACCCACTTTCATACATACCACTTTACCACATCTCCCCACTTCGCACCACCGAAAACCTAAAAAAGTGCAAAAAAAAATCATTCCAGCCAAAG |
| Bacteria-Firmicutes | *Staphylococcus aureus* | NCTC 8325 | NC_007795.1 | 1091996-1092096 | CATTTTTTTGTTTTTTAAAATAAATTTCACAAATTTTGTATAAATAGTGGTGGATAGTGGGGAGATGTGGTAAATTATATATAAGGTGAGGTGATAAAAAA |
| Bacteria-Firmicutes | *Enterococcus faecium* | SRR24 | NZ_CP038996.1 | 736432-736532 | TGAATAAAAAAAGTTGTTCCCGGATTTCCCAGAAAAGGTGGTAGAAAGTGGGGATTGTGGTAGACTAATTTTAGTTAGTGGAGGAATGGGGGGCTTTCAA |
| Bacteria-Firmicutes | *Lactococcus lactis* | L19 | CP064339.1 | 1917194-1917294 | TTGAAAGCCCCCCATTCCTCCACTAACTAAAATTAGTCTACCACAATCCCCCACTTTCTACCACCTTTTCTAGGAAATCCGGGAACAACTTTTTTTATTCA |
| Bacteria-Firmicutes | *Staphylococcus epidermidis* | ATCC 14990 | NZ_CP035288.1 | 1732100-1732200 | TTTTTTATCACCTCACCTTATTTATAATTTACCACATCTCCCCACTATCCTCCACTAATTATACAAAAAAGGTGCAAATAATTTGCTCAAACACAAAAAAC |
| Bacteria-Firmicutes | *Streptococcus pneumoniae* | SMRU 2224 | CLUP01000011.1 | 24949-25049 | CTTTCTCTCCTTAACTCACCACTATTCACCACTTCGCTCCACAACTTCATTTTACACTGTTTACCCATAAAAAAAAACCCTGAACTAGTCAGGGTTCTTTA |
| Bacteria-Actinobacteria | *Mycobacterium tuberculosis* | 401416 | CPWZ01000055.1 | 2588-2688 | TGGGGTGCTACCGCCCCCACGGCGCCCCACTCTACTCCACTCTTCCCCACCGTCAACAAGAAAAACACCCTCCGCGAGCGAATTCGGCTCACGAAACCGCA |
| Fungi-Glomeromycota | *Rhizophagus irregularis* | A1 | LLXH01006090.1 | 929-1029 | TGCTACTCACCCCACTTTAGTAAATAATGTACCACAATCCCCCACTTTGCTCCACAATTTTTAAAAAATTGTTGACAACTTTTCTGAAATTGTGTAAATAG |
| Archaea-Euryarchaeota | *Thermoplasmata archaeon* | B58_G1 | QMSW01000366.1 | 122-222 | TTTATTTCCCAAAAAATTATAAAAAAGAAATAAAAATCTTTCTTGACAAGTTAAATGGGAGATGATATACTGAAAGTGGAGTGGAGTGGGATAAAGTGGGA |
| Animalia-Arthropoda | *Abscondita terminalis* | Ate-2015 | JABVZW010003027.1 | 338757-338857 | TAATTTATCCCATTTCTGCCCACATTTACCCACTTTTTGCCACTTAATTACACTATAGGAATTAGAATTGCTTCTTGTCAAGCATTAGCTGGAGGAGCTTG |
